# ERβ regulated ovarian kisspeptin plays an important role in oocyte maturation

**DOI:** 10.1101/2020.07.27.223693

**Authors:** V. Praveen Chakravarthi, Subhra Ghosh, Katherine F. Roby, Michael W. Wolfe, M. A. Karim Rumi

**Affiliations:** Department of Pathology and Laboratory Medicine, University of Kansas Medical Center, Kansas City, Kansas, USA; Department of Anatomy and Cell Biology, University of Kansas Medical Center, Kansas City, Kansas, USA; Department of Molecular and Integrative Physiology, University of Kansas Medical Center, Kansas City, Kansas, USA; Institute for Reproduction and Perinatal Research, University of Kansas Medical Center, Kansas City, Kansas, USA

**Keywords:** Kisspeptin, kisspeptin receptor, estrogen receptor β, gonadotropin, granulosa cells, oocyte maturation

## Abstract

Kisspeptin (KISS1) signaling in the hypothalamic-pituitary (H-P) axis plays essential role in regulating gonadotropin secretion. KISS1 and KISS1 receptor (KISS1R) are also expressed in the ovary; however, the role of intraovarian KISS1 signaling remains largely unclear. Granulosa cell (GC)-specific expression of KISS1, and oocyte-specific expression of KISS1R indicate that GC-derived KISS1 may act on oocytes. Expression of KISS1 in GCs is induced by gonadotropins but it is absent in estrogen receptor β knockout (*Erβ^null^*) rats. We also observed that gonadotropin stimulation failed to induce maturation of *Erβ^null^* oocytes. Interestingly, KISS1 treatment of cumulus oocyte complexes (COCs) isolated from antral follicles promotes *in vitro* maturation of oocytes. Treatment of oocytes with KISS1 induced intracellular Ca^2+^ release, and increased activation of MAP kinase ERK1/2. KISS1 treatment also induced the expression of oocyte genes that are crucial for differentiation of GCs, and maturation of oocytes. Our findings suggest that ovarian KISS1-signaling plays an important role in gonadotropin induced follicle development and oocyte maturation.

## 1. Introduction

Estrogen receptor β (ERβ) is the predominant estrogen receptor in the ovary (Byers et al., 1997; Couse et al., 2004; Pelletier and El-Alfy, 2000; Slomczynska et al., 2001) and disruption of ERβ signaling leads ovulatory defects (Asadi et al., 2013; Lang-Muritano et al., 2018). Failure of ovulation in *Erβ^null^* mice and rats is associated with defective follicle maturation and an attenuated preovulatory gonadotropin surge (Couse et al., 2005; Rumi et al., 2017). However, treatment with exogenous gonadotropins fails to induce ovulation, suggesting a primary ovarian defect due to the loss of ERβ (Rumi et al., 2017). To identify the ERβ-regulated genes that may affect ovarian follicle development, oocyte maturation, and ovulation, we performed RNA-sequencing analyses of gonadotropin stimulated granulosa cells (GCs) and oocytes (Chakravarthi et al., 2020; Chakravarthi et al., 2019; Khristi et al., 2018). We identified *Kiss1* to be one of the most-differentially expressed gene in *Erβ^null^* GCs (Chakravarthi et al., 2020; Khristi et al., 2018). GCs showed a marked induction in *Kiss1* expression in response to gonadotropin stimulation, which was absent in *Erβ^null^* rats (Chakravarthi et al., 2020; Chakravarthi et al., 2018; Khristi et al., 2018).

Hypothalamic KISS1 regulates gonadotropin secretion, which is essential for gonadal development, onset of puberty, and induction of ovulation (Javed et al., 2015; Moore et al., 2018). In recent studies, expression of KISS1 and KISS1R in the ovaries has been detected across species (Castellano et al., 2006; Gaytan et al., 2009; Shahed and Young, 2009; Terao et al., 2004). However, the physiological role of ovarian KISS1 signaling remains largely unknown. While the expression of *Kiss1* is markedly higher in GCs, *Kiss1r* is detected predominantly in the oocytes, suggesting that GC-derived KISS1 may act on oocytes (Chakravarthi et al., 2018).

Bidirectional interactions between GCs and oocytes play vital roles in follicle development and ovulation (Craig et al., 2007; Emori and Sugiura, 2014; McNatty et al., 2003; Monniaux, 2016; Russell and Robker, 2007; Wigglesworth et al., 2013). While oocyte maturation is regulated by GC-derived factors, maturing oocytes contribute to GC differentiation and cumulus cell expansion (Craig et al., 2007; Emori and Sugiura, 2014; McNatty et al., 2003; Monniaux, 2016; Russell and Robker, 2007; Wigglesworth et al., 2013). KISS1 expressed in GCs represent the former group of signaling molecules. KISS1 expression in the ovary coincides with the preovulatory gonadotropin surge (Castellano et al., 2006) and intrafollicular KISS1 levels positively correlate with the stages of oocyte maturation (Taniguchi et al., 2017). Taken together, we hypothesize that GC-derived KISS1 may act on oocytes to effect oocyte maturation, and an attenuated expression of KISS1 in *Erβ^null^* GCs may cause failure of follicle development and oocyte maturation despite gonadotropin stimulation.

## 2. Materials and methods

### 2.1 Animal models

Wildtype and *Erβ^null^* Holtzman Sprague-Dawley (HSD) rats were studied for intraovarian kisspeptin signaling. *Erβ^null^* rat model was generated by targeted deletion of the exon 3 in the *Er*β gene (Rumi et al., 2017). Exon 3 deletion caused a frameshift and null mutation (Rumi et al., 2017). Rats were screened for the presence of mutations by PCR using tail-tip DNA samples (RED extract-N-Amp Tissue PCR Kit, Sigma-Aldrich) as described previously (Rumi et al., 2014; Rumi et al., 2017). All procedures were performed in accordance with the protocols approved by the University of Kansas Medical Center Animal Care and Use Committee.

### 2.2 Gonadotropin treatment

4-wk-old *Erβ^null^* and age-matched wildtype female rats were used to evaluate gonadotropin-induced follicular development. Synchronized follicular growth was performed by intraperitoneal injection of 30 IU pregnant mare serum gonadotropin (PMSG; BioVendor LLC, Asheville, NC). Forty-eight hours after the PMSG injection, 30 IU of human chorionic gonadotropin (hCG; BioVendor) was injected intraperitoneally. Rats were sacrificed prior to gonadotropin treatment (Basal), 48h after PMSG treatment (PMSG), 4h and 24h after hCG injection to PMSG treated rats and their ovaries were collected (**Fig. 1B**) as described previously (Chakravarthi et al., 2020; Chakravarthi et al., 2019; Chakravarthi et al., 2018).

**Fig. 1.**
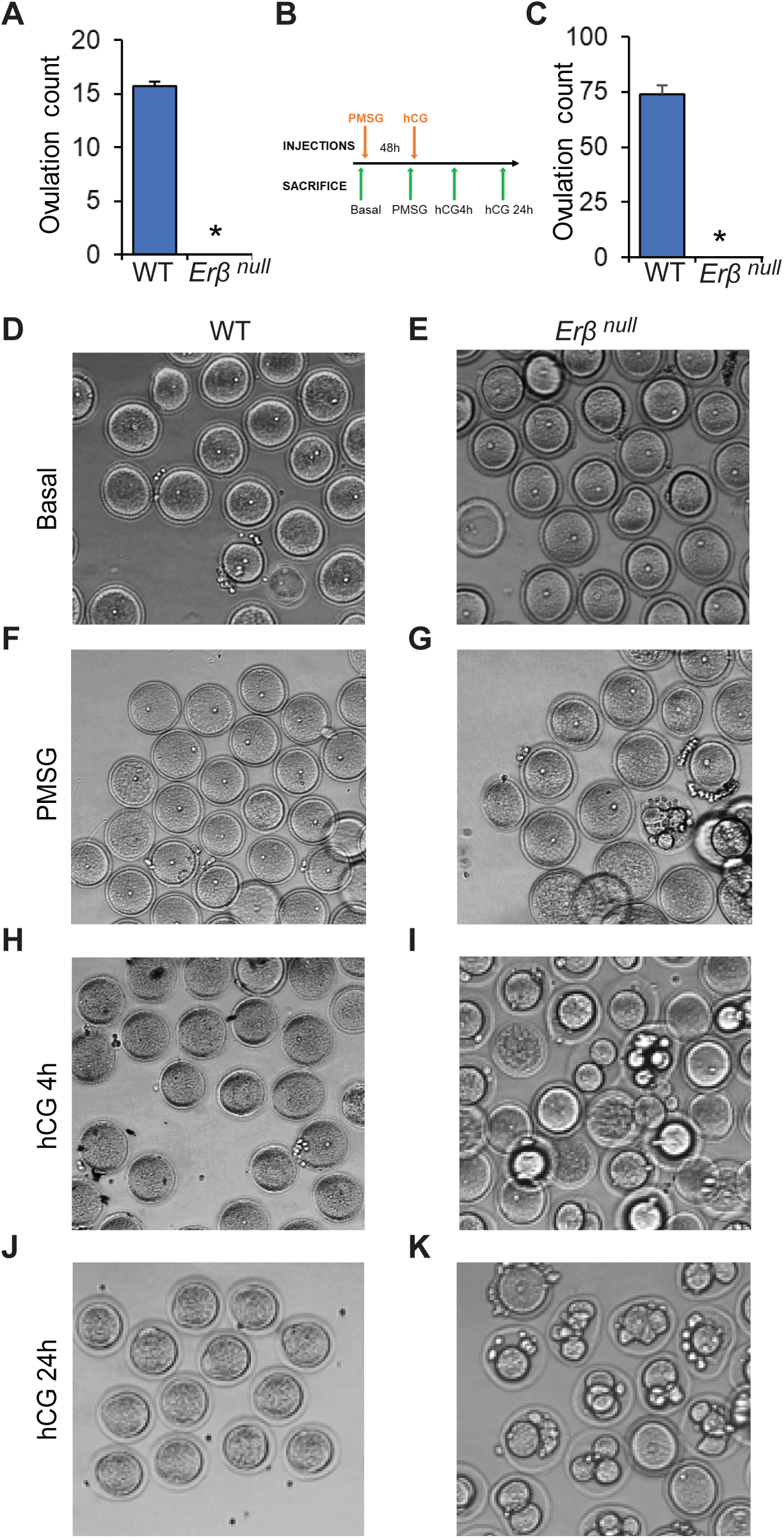
Abnormal oocyte maturation in *Erβ^null^* rats. 8-12 wk-old *Erβ^null^* adult rats are infertile and they failed to ovulate either during natural estrus cycle (**A**) or induced by exogenous gonadotropins to 4 wk-old rats (**C**). Oocytes isolated from 4-wk-old wildtype (WT) and *Erβ^null^* rat ovaries at different stages of gonadotropin treatment were examined after removal of cumulus cells. Although unstimulated (Basal **D** and **E**) oocytes of both genotypes appeared similar, abnormal division and fragmentation was observed in *Erβ^null^* oocytes after PMSG 48h (**F** and **G**), 4h after hCG (**H** and **I**), and 24h after hCG (**J** and **K**) treatment.

### 2.3 Isolation of oocytes and GCs

Cumulus oocyte complexes (COCs) were isolated from the large antral follicles by needle puncture under microscopic observation as described previously (Chakravarthi et al., 2020; Chakravarthi et al., 2019; Chakravarthi et al., 2018). All cumulus cells were removed from the oocytes by mechanical pipetting followed by repeated washings into fresh media 199 using capillary suctions. GCs were collected by filtration of the remaining cells through 40μM cell strainers (Fisher Scientific).

### 2.4 RNA-sequencing of GCs and oocytes

Oocytes and GCs were collected from 4-wk-old wildtype and *Erβ^null^* rats with or without gonadotropin treatment. Total RNA was extracted using TRI Reagent (Millipore-Sigma) following the manufactures instructions. Approximately 500ng of total RNA was used for the RNA-seq library preparation. Libraries were prepared by using a TruSeq standard mRNA kit (Illumina) following manufacturer’s instructions. The cDNA libraries were sequenced at the Molecular Biology Core Laboratory of Mayo Clinic (Rochester, MN). RNA-Seq data were analyzed using CLC Genomics Workbench (Qiagen Bioinformatics). Gene expression values were reported as TPM (Transcripts per million base pairs). All the RNA seq data are available at SRA (SRX6376730, SRX6376729, SRX6376735, SRX6376752, SRX6376751, SRX6376757, SX6761695, SRX6761696, SRX6761697, SRX6761698, SRX6761699 and SRX67617) and the data articles are published in Data in Brief journal (Chakravarthi et al., 2020; Chakravarthi et al., 2019).

### 2.5 In vitro maturation of rat oocytes

COCs were collected from the antral follicles of 4-wk-old wildtype and *Erβ^null^* rats 48h after PMSG treatment. After washing three times in *in vitro* maturation media (BO-IVM, IVF Biosciences, UK) the COCs were placed in 20 µl drops of IVM medium in 35-mm culture dishes, covered with light weight mineral oil, and incubated at 37°C in a humidified CO2 incubator for an additional 24h (Kona et al., 2016; Praveen Chakravarthi et al., 2015). At the end of incubation, oocytes were denuded of the cumulus cells by repeated pipetting through a fine bore glass pipette and examined microscopically for extrusion of the polar body as indicative of oocyte maturation to MII stage (Kona et al., 2016; Praveen Chakravarthi et al., 2015).

### 2.6 Calcium release assay

4-wk-old wildtype rats were administered with PMSG, and 48h later COCs were isolated. COCs were incubated with 5µM Fluo-3AM (ThermoFisher Scientific) solution containing 20% Pluronic F-127 (ThermoFisher Scientific) for 30 min in a 37°C incubator. All cumulus cells were removed from the oocytes by pipetting followed by repeated washings into fresh media using capillary suction and oocytes were placed into HBSS drops containing 1nM of rat kisspeptin-10 (KP-10) (Tocris Bioscience, Bristol, UK). Oocytes were monitored under a fluorescent microscope for 3 min and images were recorded every 30 sec.

### 2.7 Protein extraction and western blotting

COCs were collected from 4-wk-old wildtype rats 48h after PMSG administration. Oocytes and GCs were purified as described in Section 2.2 and proteins were extracted in 1X SDS lysis buffer (62.5mMTris-HCl pH 6.8, 2% SDS, 42 mM dithiothreitol, 10% glycerol, and 0.01% bromophenol blue), containing protease and phosphatase inhibitors (Cell Signaling Technologies, Danvers, MA). In addition, oocytes were cultured with or without 1nM or 10nM KP-10. After treatment, oocytes were denuded mechanically, proteins were extracted in 1XSDS buffer. Oocyte or GC cell-lysates in 1XSDS buffer were sonicated to shear DNA and reduce viscosity, heat denatured, and separated on a 4-20% SDS-PAGE. Electrophoresed proteins were transferred from the gel to PVDF membranes, blocked with 5% skim milk in TBST (1XTBS buffer containing 0.1% Tween-20), and incubated overnight at 4°C with specific primary antibodies at the appropriate dilution in blocking solution (**Supplementary Table 1**). After removing the unbound primary antibody solution, membranes were washed with TBST, blocked, and incubated with peroxidase conjugated anti-mouse, or anti-rabbit secondary antibodies (Jackson Immunoresearch, West Grove, PA) at a dilution of 1:10,000 to 50,000, and the immunoreactivite signals were visualized with Luminata Crescendo HRP substrate (Millipore Sigma, Burlington, MA).

### 2.8 Preparation cDNA and RT-qPCR analyses

Total RNA was extracted from the GCs and oocytes using TRI Reagents (Sigma-Aldrich). 1000 ng of total RNA from each sample was used for the preparation of cDNAs using High-Capacity cDNA Reverse Transcription Kits (Applied Biosystems, Foster City, CA). RT-qPCR amplification of cDNAs was carried out in a 10µl reaction mixture containing Applied Biosystems Power SYBR Green PCR Master Mix (Thermo Fisher Scientific). Amplification and fluorescence detection of RT-qPCR were carried out on Applied Biosystems Quant Studio Flex 7 Real Time PCR System (Thermo Fisher Scientific). The ΔΔCT method was used for relative quantification of target mRNA expression level normalized to Rn18s (18S rRNA). A list of qPCR primers is shown in **Supplementary Table 2**.

### 2.9 Statistical analyses

Ovulation counting was performed on 6 individual rats of the same genotype at each time point. Each RNA-sequencing library was prepared using pooled RNA samples form three individual wildtype or *Erβ^null^* rats and each group of RNA-sequencing consisted of three libraries. Gene expression analyses were performed on at least 6 individual rats. The experimental results are expressed as mean ± SE. Statistical comparisons between two means were determined with Student’s t-test while comparisons among multiple means were evaluated with ANOVA and the significance of mean differences was determined by Duncan post hoc test, with P ≤ 0.05 considered a significant level of difference. All statistical calculations were done with SPSS 22 (IBM, Armonk, NY).

## 3. Results

### 3.1 ERβ is essential for oocyte maturation

8-12 wk old adult *Erβ^null^* rats failed to ovulate during a natural estrous cycle (**Fig. 1A**). Administration of exogenous gonadotropins also failed to induce ovulation in 4-wk-old *Erβ^null^* rats (**Fig. 1C**). We observed that *Erβ^null^* oocytes isolated from the antral follicles prior to gonadotropin stimulation appeared similar to those isolated from wildtype follicles (**Fig. 1D** and **E**). However, gonadotropin treatment resulted in an abnormal maturation of the *Erβ^null^* oocytes (**Fig. 1F-K**). Development of abnormal and fragmented nuclei in *Erβ^null^* oocytes became more evident after hCG treatment (**Fig.1H-K**).

### 3.2 Kiss1 is expressed in GCs and Kiss1r in oocytes

We observed a prominent expression of *Kiss1* in gonadotropin treated GCs (**Fig. 2A**) while *Kiss1r* in oocytes (**Fig. 2B**). RNA-sequencing revealed a high level of *Kiss1* transcripts in wildtype GCs, but the expression was severely reduced or absent in *Erβ^null^* GCs (**Fig. 2C**). These findings were validated by qRT-PCR analyses (**Fig. 2D**). In contrast to *Kiss1, Kiss1r* expression did not show any significant difference between the wildtype or *Erβ^null^* oocytes (**Fig. 2E** and **F**).

**Fig. 2.**
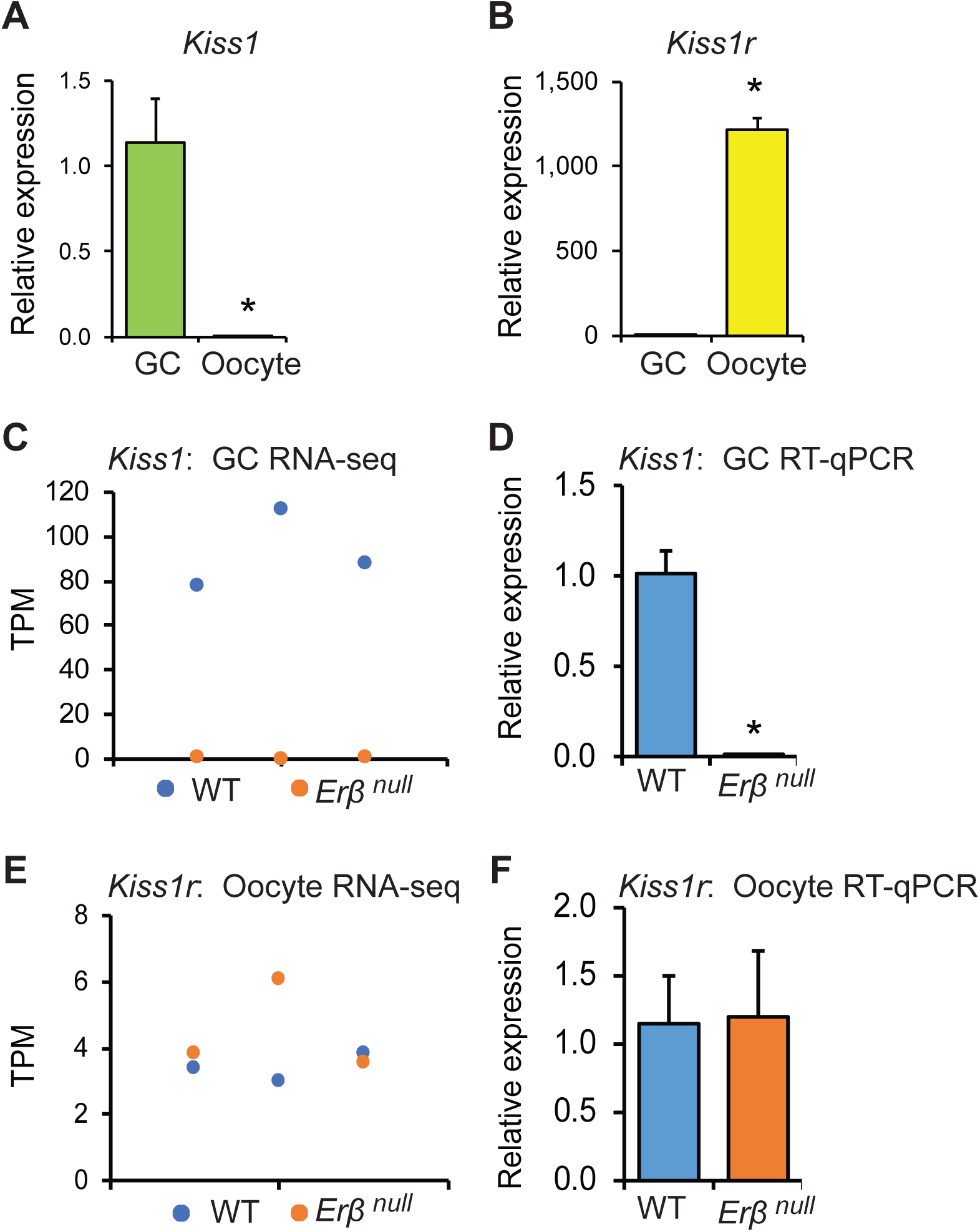
*Kiss1* and *Kiss1r* expression in rat ovary. Oocytes and granulosa cells (GCs) were isolated from the ovaries collected 4h after hCG administration to PMSG treated 4-wk-old rats. RT-qPCR analyses showed that expression of *Kiss1* was remarkably higher in (GCs) (**A**), whereas *Kiss1r* in the oocytes (**B**). RNA sequencing analyses showed a significantly higher expression of *Kiss1* in wildtype (WT) GCs compared to *Erβ^null^* GCs (**C**), but a similar level of *Kiss1r* expression in both WT and *Erβ^null^* oocytes (**E**). RT-qPCR data from GCs and oocytes further confirmed the results of RNA-sequencing (**D** and **F**). RNA-sequencing data are presented in triplicates. RT-qPCR data are represented as mean ± SE. n = 6. *P ≤ 0.05.

### 3.3 KISS1 treatment induces in vitro maturation of oocytes

We observed that *Erβ^null^* oocytes exhibit defects in *in vitro* maturation. While approximately 55% of wildtype oocytes mature to MII stage (**Fig. 3A**), less than 35% of *Erβ^null^* oocytes undergo *in vitro* maturation (**Fig. 3D**). Remarkably, treatment with rat kisspeptin 10 (KP-10) increased the rate of oocyte maturation in both wildtype (**Fig. 3B** and **C**) and *Erβ^null^* oocytes (**Fig. 3E** and **F**).

**Fig. 3.**
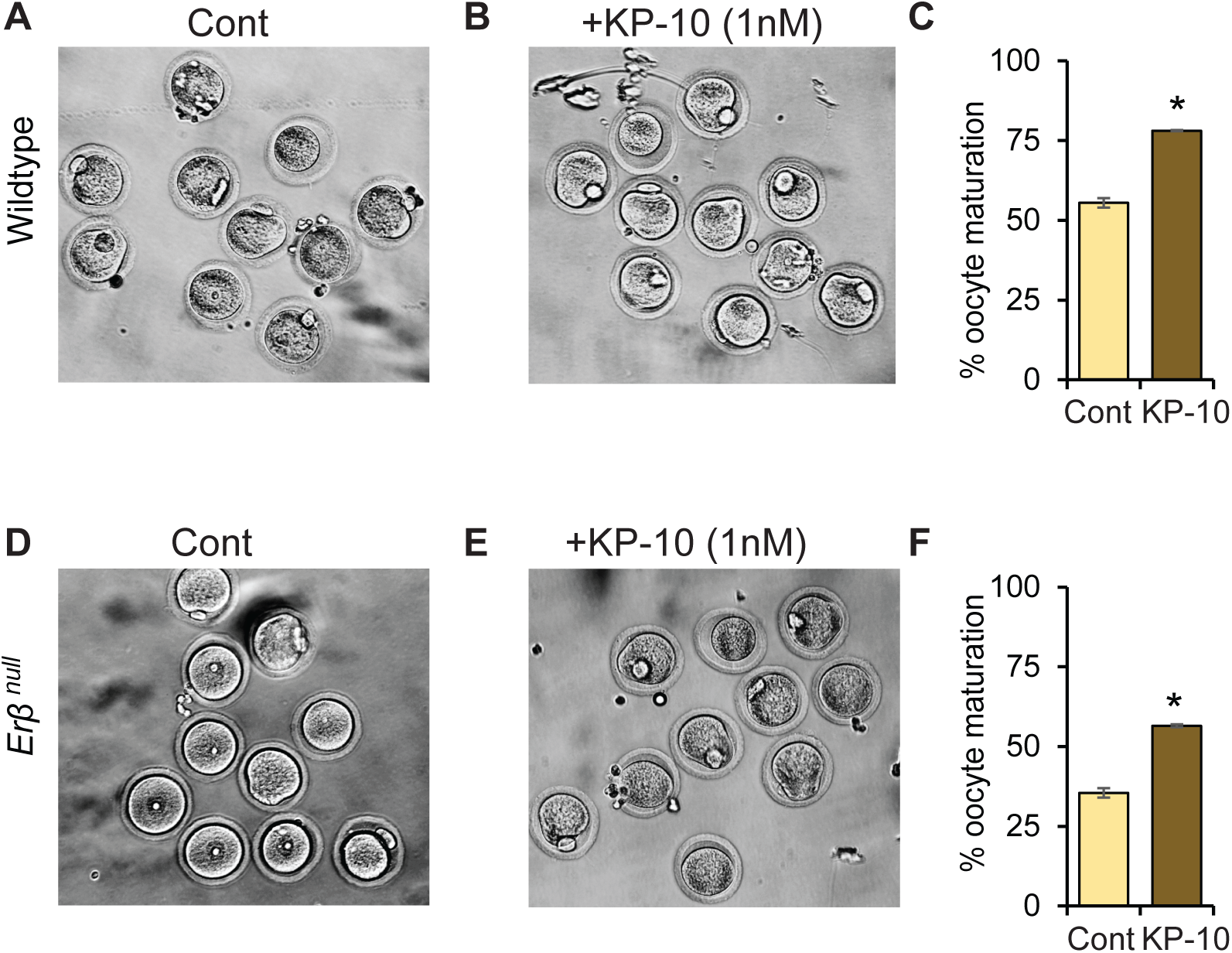
Role of KISS1 in oocyte maturation. 48h after PMSG treatment, cumulus oocyte complex (COCs) were isolated from 4-wk-old wildtype and *Erβ^null^* rats and cultured in BO-IVM in presence or absence of 1nM of rat kisspeptin 10 (KP-10) for 20h. Approximately 55% of wildtype (**A** and **C**) and only 35% of *Erβ^null^* oocytes achieved meiotic maturation (MII) in BO-IVM culture (control) (**D** and **F**). Remarkably, addition of KP-10 to BO-IVM increased the *in vitro* maturation to 78% of wildtype (**B** and **C**) and 56% of *Erβ^null^* oocytes (**E** and **F**). Each treatment group contained at least 25 oocytes at each time point and repeated three times. Cont, Control.

### 3.4 KISS1 signaling in rat oocytes

Western blot analyses detected the expression of KISS1R in oocytes but not in GCs (**Fig. 4A**). Treatment of oocytes with rat KP-10 induced a prominent intracellular Ca^2+^ release (**Fig. 4B-E**). KP-10 treatment also activated the ERK1/2 in oocytes (**Fig. 4F**), but not the AKT (**Fig. 4G**). While 1nM of KP-10 induced marked activation of ERK1/2, an increased concentration failed to show such an effect.

**Fig. 4.**
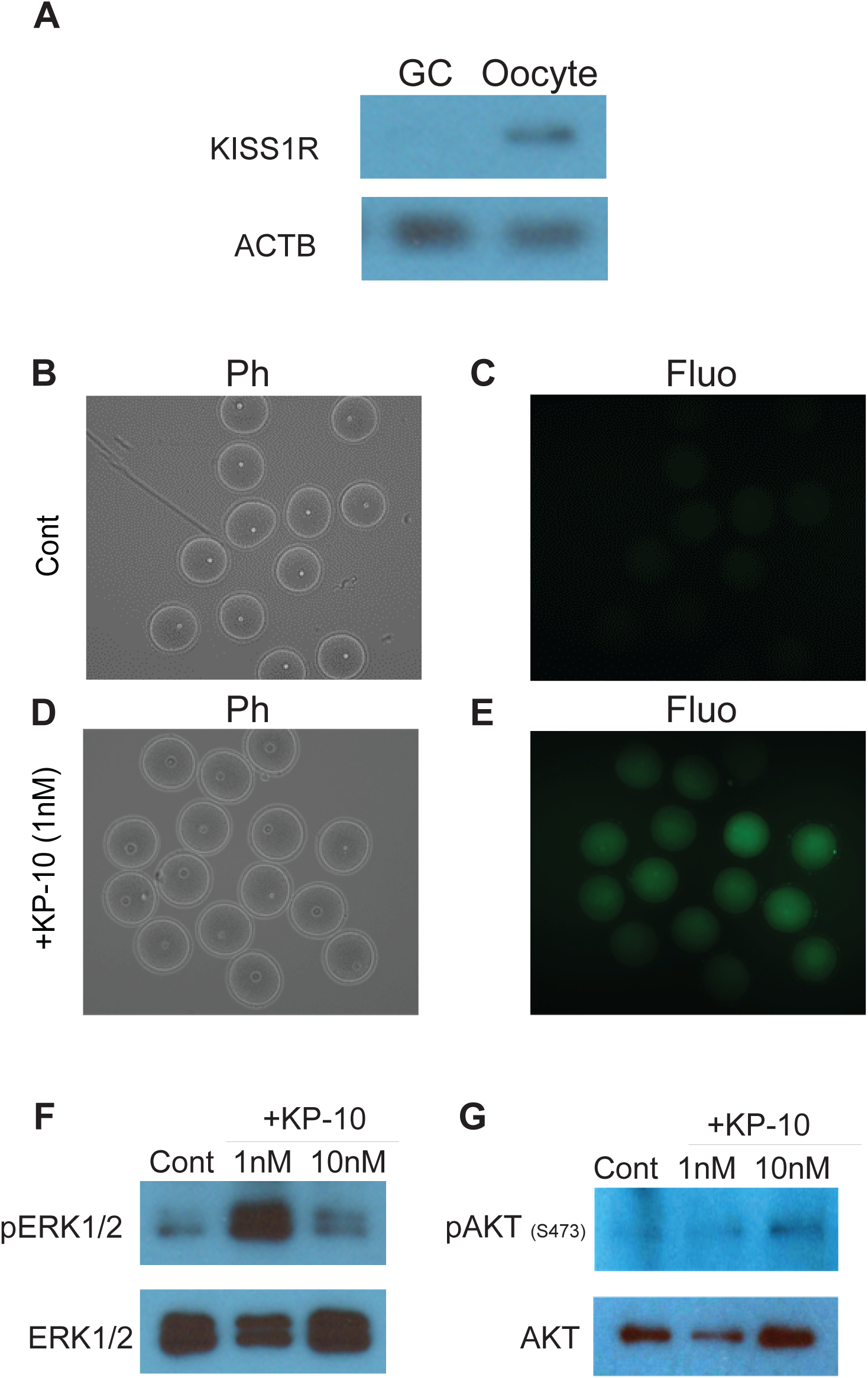
Role of KISS1 in inducing Calcium efflux and ERK signaling. The expression of KISS1R was significantly higher in oocytes compared to granulosa cells (GCs) as shown by western blot analysis (**A**). Oocytes from PMSG treated rats were examined for calcium efflux using a Fluo-3 assay kit. Calcium efflux increased after the treatment with rat kisspeptin 10 (KP-10) (**E**) compared to control (**C**). Treatment with 1nM of KP-10 also markedly elevated the phosphorylation of ERK1/2 (**F**) but not AKT (**G**) in oocytes. Ph, Phase contrast; Fluo, Fluorescence microscopy; Cont, Control.

### 3.5 KP-10 treatment induces genes involved in oocyte maturation

Expression of genes known to be involved in regulating oocyte maturation were assessed by RT-qPCR analyses (**Fig. 5**). KP-10 treatment upregulated the expression of *Gdf9* and *Bmp15*, (**Fig. 5 A** and **B)**, whereas it downregulated the expression of *Bmp7* (**Fig. 5 C**). Expression of genes involved in protein kinase activation required for oocyte maturation including *cMos, Kit* and *Aurka* (**Fig. 5 D-F)** were also increased by KP-10 stimulation of oocytes. Similarly, KP-10 treatment increased the expression of genes involved in meiosis including *Wee2, Cdc25b* and *Rec8* (**Fig. 5 G-I**). RNA binding proteins like *Dazl, Btg4* and *Papbc1l*, which are known to induce mRNA stability and selective activation of genes were also upregulated with KP-10 treatment (**Fig. 5 J-L**).

**Fig. 5.**
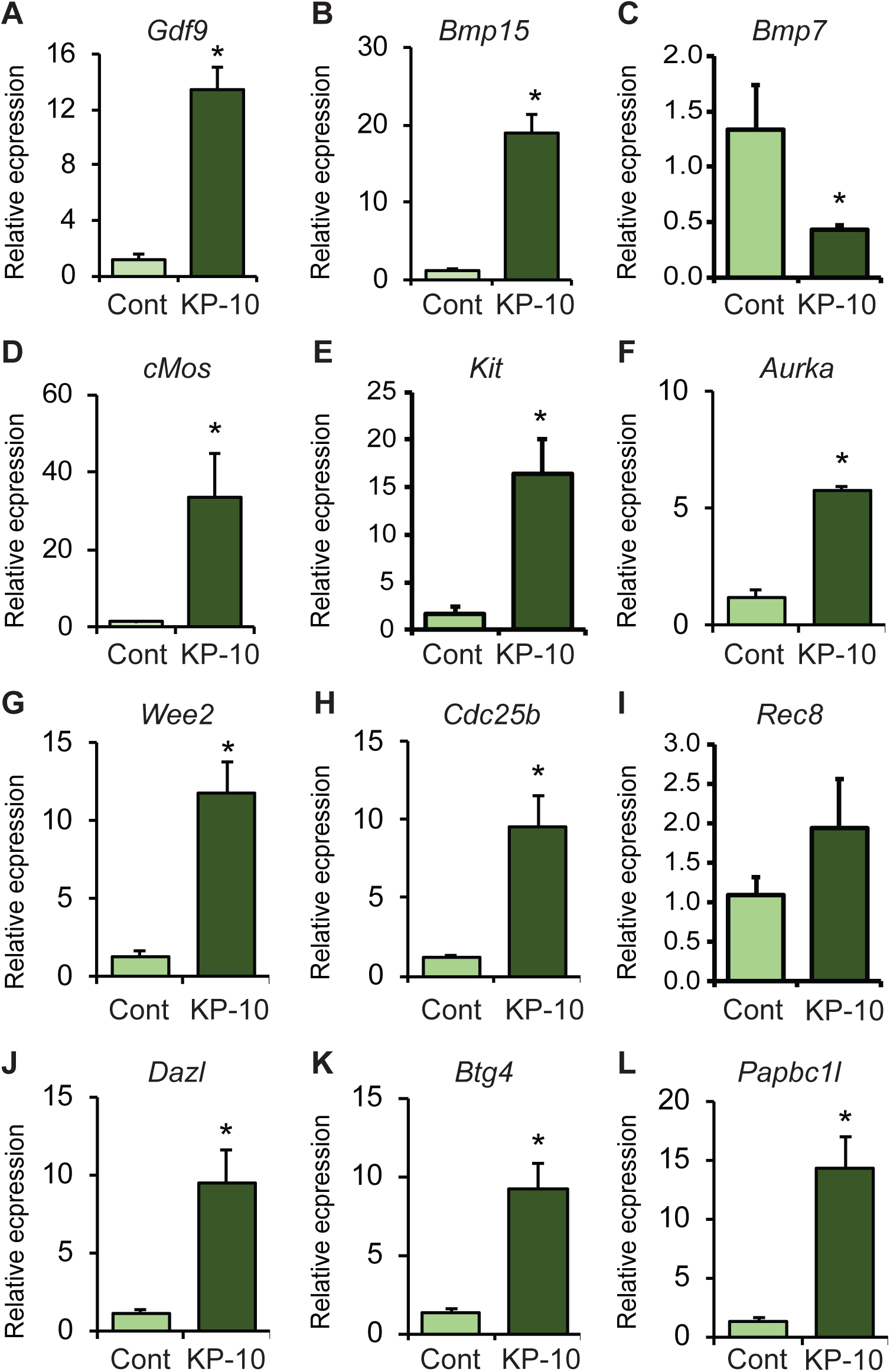
KP-10 treatment regulated genes involved in inducing oocyte maturation. 48h after PMSG treatment, cumulus oocyte complex (COCs) were isolated from 4-wk-old wildtype female rats and treated with 1nM of rat kisspeptin 10 (KP-10) for 30 min. KP-10 treatment upregulated the mRNA levels of *Gdf9, Bmp15*, but downregulated *Bmp7* (**A-C**). It also upregulated the mRNA levels of protein kinases *cMos, Kit, and Aurka* (**D-F**), as well as *Wee2* and *Cdc25b* (**G** and **H**) important for meiotic maturation. Moreover, KP-10 treatment upregulated the mRNA levels of RNA binding proteins *Dazl, Btg4*, and *Papbc1l* (**J-L**) required for selective mRNA stability during oocyte maturation. RT-qPCR data represent the mean ± SE. *P < 0.05, n = 6.

## 4. Discussion

We have studied the role of ovarian derived KISS1 using an *Erβ^null^* mutant rat model that fails to induce *Kiss1* gene expression in GCs in response to gonadotropins (Chakravarthi et al., 2020; Chakravarthi et al., 2018; Khristi et al., 2018) and suffer from defective oocyte maturation. Intraovarian *Kiss1* gene expression is induced by the preovulatory gonadotropin surge that initiates the final steps of follicle development and oocyte maturation (Castellano et al., 2006). We also detected a dramatic upregulation of *Kiss1* gene expression in GCs following the administration of hCG to PMSG primed rats (Chakravarthi et al., 2020; Chakravarthi et al., 2018; Khristi et al., 2018). The expression pattern of *Kiss1* and *Kiss1r* (Chakravarthi et al., 2018) suggest that GC-derived KISS1 may act on the KISS1R in oocytes to regulate oocyte maturation (**Fig. 6**). An absence of *Kiss1* gene induction in *Erβ^null^* GCs may be linked to the defective oocyte maturation in *Erβ^null^* rats in response to gonadotropin treatment.

**Fig. 6.**
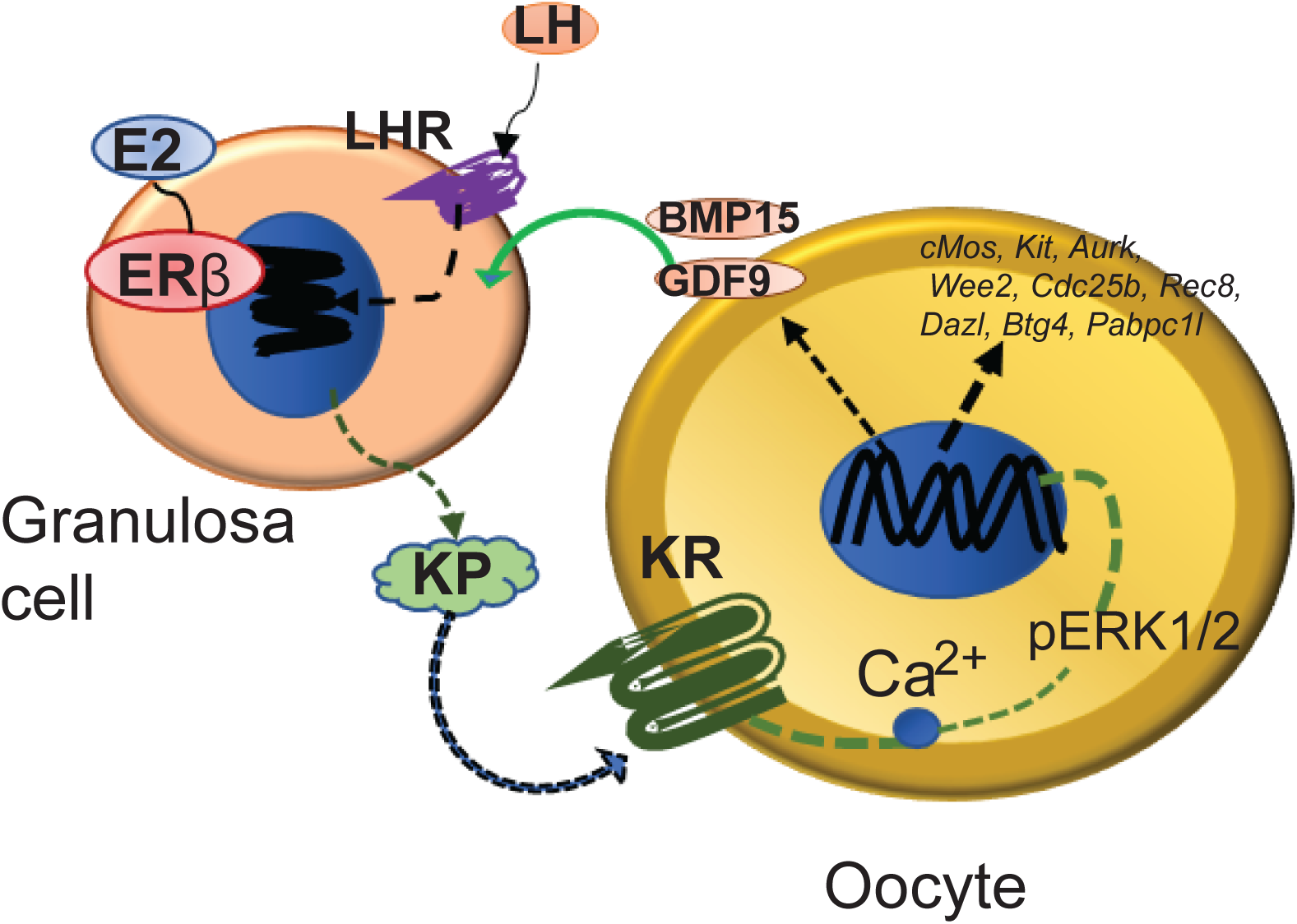
Intraovarian role of ovarian derived KISS1. Based on our findings, we conclude that the preovulatory LH surge induces the expression of KISS1 (KP) by granulosa cells (GCs), which is ERβ-dependent. GC-derived KP acts on the KISS1R (KR) expressed in oocytes to increase Ca2+ release, ERK phosphorylation, and modulation of genes, which are important for inducing oocyte maturation.

We observed that treatment of oocytes with a low dose (1nM) of KP-10 improved *in vitro* maturation of rat oocytes. Previous studies on pig (Saadeldin et al., 2012) and sheep (Byri et al., 2017) oocytes also support our findings. Although *Erβ^null^* oocytes showed a reduced rate of *in vitro* maturation, both of wildtype and *Erβ^null^* oocytes responded to rat KP-10 treatment showing a similar improvement. This can be explained by similar levels of *Kiss1r* expression in wildtype and *Erβ^null^* oocytes (Fig. 2E, F) and we would assert normal intracellular signaling induced by KP-10.

KISS1R protein is expressed in rat oocytes and responds to KISS1 treatment. Intracellular Ca^2+^ release immediately after KP-10 treatment further verifies the presence of KISS1R and indicates specific KISS1/KISS1R signaling. KISS1R is a G-protein coupled receptor that signals through intracellular Ca^2+^ release. Activation of MAPK kinase ERK1/2 following kisspeptin treatment has been reported with KISS1/KISS1R signaling in hypothalamic neurons (d’Anglemont de Tassigny and Colledge, 2010) and we have confirmed that this signaling pathway is maintained in oocytes. Conversely, we did not observe activation of AKT by KP-10 in oocytes, but this is a typical response to KISS1/KISS1R signaling in the hypothalamic neurons (d’Anglemont de Tassigny and Colledge, 2010). In oocytes, Ca^2+^ stored in the endoplasmic reticulum and generates a Ca^2+^ efflux that is important for signaling and inducing oocyte maturation (Kline, 2000; Stricker et al., 1998; Tosti, 2006; Wang and Machaty, 2013). Ca^2+^ efflux has also been reported to induce RAF/MEK/ERK pathway through activation of RAS (Atherfold et al., 1999; Li et al., 2005). Furthermore, ERK1/2 activation has also been found essential for oocyte maturation (Choi et al., 1996). We observed that KP-10 induced Ca^2+^ efflux in oocytes, which resulted in an upregulation of ERK1/2 activation. Thus KP-10 induction of both Ca^2+^ efflux and ERK1/2 activation may be responsible for the enhanced oocyte maturation that was observed in the current study.

To elucidate the mechanisms downstream of Ca^2+^ signaling and ERK1/2 activation, we investigated transcript levels of genes that play important roles in oocyte maturation. We observed that KP-10 treatment upregulated the expression of *Gdf9, Bmp15, cMos, Kit, Aurka, Wee2, Cdc25b, Dazl, Btg4*, and *Papbc1l*. BMP15 and GDF9 are known to promote cumulus expansion and regulate oocyte maturation (Chang et al., 2016; McNatty et al., 2005a; McNatty et al., 2005b; Persani et al., 2014). ERK/MAPK pathway has been shown to induce the expression of *Bmp15* and *Gdf9* which are essential for growth and differentiation of GCs during oocyte maturation (Mottershead et al., 2012; Reader et al., 2011). KP-10 treatment reduced the expression of BMP7 which has been shown to inhibit oocyte maturation (Yang et al., 2018). cMOS has been reported to be involved in the spindle formation, activation of maturation promoting factor, and oocyte maturation (Araki et al., 1996; Chang et al., 2016; Choi et al., 1996; Hashiba et al., 2001). KIT/KITL signaling also enhances the oocyte maturation in *in vitro* by overcoming the inhibitory effect of NPPC signaling (Medvedev et al., 2011; Ye et al., 2009). AURKA is a serine /threonine kinase that accumulates on microtubule organizing centers and is involved in the resumption of meiosis (Saskova et al., 2008). WEE2 is involved in the inhibitory phosphorylation of CDC2 that drives the exit from metaphase II (MII), promoting oocyte maturation (Oh et al., 2011). CDC25 is a dual specificity phosphatase that activates cyclin dependent kinases and regulates meiotic maturation of oocytes (Lincoln et al., 2002). *Dazl* encodes an RNA binding protein that plays an important role in spindle assembly, MI-MII transition and oocyte maturation (Chen et al., 2011). PABPC, a poly A binding protein that increases stability of selective mRNAs, promotes mRNA translation and improves oocyte maturation (Lowther and Mehlmann, 2015). In addition, BTG4 is involved in selective degradation of oocyte mRNA that is required for maternal-zygotic transition (Yu et al., 2016). We speculate that these KP-10 induced changes in gene expression play important role in promoting oocyte maturation.

It is well established that KISS1 regulation of oocyte maturation and induction of ovulation is mediated indirectly through gonadotropin secretion from the H-P axis. Our findings indicate that the role for KISS1 in regulating female fertility can be explained beyond the HP-axis to intrafollicular expression of KISS1 and signaling, which plays an important role in regulating oocyte maturation.

## Acknowledgements

This study was supported in part by the grant supports from KUMC SOM, COBRE and K-INBRE.

