## Supplemental Table 1 for "ERβ regulated ovarian kisspeptin plays an important role in oocyte maturation"

**Table 1. List of antibodies used in the western blot assays**

| **Target** | **Name of Antibody** | **Manufacturer, Catalog number** | **Species, Mono or Polyclonal** | **Primary**  **Dilution** | **2ndary**  **Dilution** |
| --- | --- | --- | --- | --- | --- |
| KISS1R | Anti-KISS1R | MyBioSource, MBS8501891 | Rabbit; polyclonal | 1,000 | 10,000 |
| ACTB | Anti-*β*-actin | Sigma-Aldrich (AC15), A 1978 | Mouse; monoclonal | 25,000 | 50,000 |
| pAKT | Anti-Phospho-Akt (Ser473) | Cell Signaling Technology, 4060 | Rabbit; monoclonal | 2000 | 25,000 |
| AKT | Anti- Akt | Cell Signaling Technology, 4691 | Rabbit; monoclonal | 2000 | 25,000 |
| pERK | Anti-Phospho-ERK1-T202/Y204 + ERK2-T185/Y187 | Cell Signaling Technology, 4370 | Rabbit; monoclonal | 2000 | 25,000 |
| ERK | Anti-p44/42 MAPK (ERK1/2) | Cell Signaling Technology, 4695 | Rabbit; monoclonal | 1000 | 25,000 |
