## Supplemental Table 2 for "ERβ regulated ovarian kisspeptin plays an important role in oocyte maturation"

**Table 1. List of primers used in the qPCR assays**

| **Symbol** | **Reference mRNA** | **Forward Primer** | **Reverse Primer** | **Amplicon** |
| --- | --- | --- | --- | --- |
| *Kiss1* | NM_181692.1 | 26F-TGCTGCTTCTCCTCTGTGTG | 174R-AGGCTTGCTCTCTGCATACC | 149bp |
| *Kiss1r* | NM_001301151.1 | 502F-CTGGGAGACTTCATGTGCAA | 614R-GGGAACACAGTCACGTACCA | 113bp |
| *Gdf9* | NM_021672.1 | 330F-CTACAATACCGTCCGGCTCT | 475R-GCAAGACCGATTTGAGTAAGTG | 146bp |
| *Bmp15* | NM_021670.1 | 525F-GAGGCTGATAAAGCCGTCAG | 663R-AAGTTGATGGCGGTAGACCA | 139bp |
| *Bmp7* | NM_001191856.2 | 1080F-AGGAGGGCTGGTTGGTATTT | 1366R-CATCCTCAGTGCCTCTTGGT | 287bp |
| *cMos* | NM_020102.4 | 453F-ACCAGGAACCTACGTGCATC | 532R-CGGACTATGTTGTCGTGGTG | 80bp |
| *Kit* | NM_022264 | 2769F-GATGCTGATCCCCTGAAAAG | 2962R-AGGAGAGGCTGTGTGGAAGA | 194bp |
| *Aurka* | NM_153296.2 | 521F-AGCTCCGGAGAGAAGTGGA | 713R-GCCAACTCCGTGATGTAGGT | 193bp |
| *Wee2* | XM_008762866 | 4109F-ATCAAGAGGCTGGATGGATG | 4218R-CCAAGCACTGCATGAGCATA | 109bp |
| *Cdc25b* | XM_006243702.3 | 521F-TGAGTGCTCCCTGTCATCTG | 711R-TATGTCCAGCAGCCTCACTG | 190bp |
| *Rec8* | NM_001011916 | 119F- TCTTGGAGCTTCGCATTTCT | 317R- TTCAGGTATTCCCGCTTCAC | 199bp |
| *Dazl* | NM_001109414 | 153F-CTGTGCCACCTTCGAGTTTT | 278R-CTGGTGGTTGCTGATGAAGA | 125bp |
| *Btg4* | NM_001013176 | 452F-TGGGACTTCCAAAGGAGATG | 575R-ACCATCTGCCAAATGCTTTC | 124bp |
| *Pabpc1l* | XM_008775624 | 126F-CGGCACTATTCTTTCCATCC | 291R-TCTGTGGGACCACATGATTC | 166bp |
